# Balancing Act: Groundwater microbiome’s resilience and vulnerability to hydroclimatic extremes

**DOI:** 10.1101/2025.02.14.638306

**Authors:** He Wang, Martina Herrmann, Simon A. Schroeter, Christian Zerfaß, Robert Lehmann, Katharina Lehmann, Arina Ivanova, Georg Pohnert, Gerd Gleixner, Susan E. Trumbore, Kai Uwe Totsche, Kirsten Küsel

**Author notes:** **Corresponding author:** Kirsten Küsel.

## Abstract

Groundwater health is increasingly threatened by climate change, which alters precipitation patterns, leading to groundwater recharge shifts. These shifts impact subsurface microbial communities, crucial for maintaining ecosystem functions. In this decade-long study of carbonate aquifers, we analyzed 815 bacterial 16S rRNA gene datasets, 226 dissolved organic matter (DOM) profiles, 387 metabolomic datasets, and 174 seepage microbiome sequences. Our findings reveal distinct short- and long-term temporal patterns of groundwater microbiomes driven by environmental fluctuations. Microbiomes of hydrologically connected aquifers exhibit lower temporal stability due to stochastic processes and greater susceptibility to surface disturbances, yet they demonstrate remarkable resilience. Conversely, isolated aquifer microbiomes show resistance to short-term changes, governed by deterministic processes, but exhibit reduced stability under prolonged stress. Variability in seepage-associated microorganisms, DOM, and metabolic diversity further drive microbiome dynamics. These findings highlight the dual vulnerability of groundwater systems to acute and chronic pressures, emphasizing the critical need for sustainable management strategies to mitigate the impacts of hydroclimatic extremes.

## Introduction

Groundwater, the largest global reservoir of accessible freshwater, is increasingly jeopardized by the dual pressures of anthropogenic activities and climate change, with widespread implications for water availability, quality, and ecosystem health^1,2,3^. Future climate scenarios predict profound shifts in groundwater recharge, driven predominantly by changing precipitation patterns and the intensification of hydroclimatic extremes^2,3^. These shifts threaten groundwater sustainability, particularly in vulnerable carbonate aquifers where water storage and flow dynamics are intricately linked to lithology and hydrological connectivity^4,5^. Understanding the resilience of groundwater ecosystems and their ability to maintain ecological functions under such pressures is critical for sustainable management strategies^3,6^.

Microbial communities are central to groundwater ecosystem health, playing pivotal roles in biogeochemical cycling, contaminant degradation, and ecosystem resilience^7,8,9^. Traditionally considered static due to the relatively constant conditions of subsurface environments, this view has persisted largely due to the lack of long-term observational studies. More recent studies challenge this notion by demonstrating that groundwater microbiomes exhibit a dynamic nature, driven by both short-term recharge events^10,11,12^ and longer-term environmental changes^13^. For instance, microbial immigration during recharge can trigger compositional shifts^12,13^, while prolonged droughts may alter microbial activity through changes in water chemistry^14^. Additionally, rain infiltration can mobilize soil microorganisms, transporting them to the groundwater via seepage^15,16,17,18^. These findings underscore the importance of hydrological connectivity as a key factor influencing microbial dispersal, assembly, and ecosystem resilience^15,19,20,21^.

Hydrological connectivity, shaped by aquifer permeability, flow dynamics, and recharge patterns, governs the flux of surface-derived inputs into groundwater and mediates microbial dispersal^5,15,19,22^. Hydrologically connected aquifers, often characterized by karstification and higher permeability, are particularly susceptible to surface-derived disturbances such as nutrient influxes, organic contaminants, and pathogens^4,22,23^. In contrast, hydrologically isolated systems may exhibit greater stability^11^ but face significant stress under conditions of prolonged environmental change, such as reduced recharge during droughts^14^,. These contrasting dynamics necessitate a framework for quantifying temporal stability (consistency), resistance (insensitivity to disturbance), and resilience (ability to recovery) of groundwater microbiomes across gradients of hydrological connectivity^9,21^.

While previous studies have focused on microbiome compositional stability across ecosystems, functional stability is also a key indicator of ecosystem health^13,24,25,26^. To this end, we applied metabolomics techniques to probe system activity in our long-term study^27^. Untargeted liquid chromatography–high-resolution mass spectrometry (LC-HRMS) can profile environmental meta-metabolomes, comprising metabolites released by microorganisms and anthropogenic activity. This approach is complemented by direct infusion–high-resolution mass spectrometry (DI-HRMS) resolving the full complexity of DOM without chromatographic bias^28,29,30,31^. This integrated information from both approaches together with microbiome function analysis using PICRUSt2 resolves quantitatively the impact of the organic chemical landscape on microbiome diversity^32,33^. This allowed us to explore groundwater microbiome stability, variability, and overall ecosystem health.

Our work expands directly on discussions highlighting not only the impacts of climate change on groundwater recharge^2^, but also on groundwater quality^14^. Building on these foundations, we investigate the critical links between hydrological connectivity and groundwater microbiome stability in carbonate aquifers. Here, we conducted a groundwater study along a hillslope well transect in the Hainich Critical Zone Exploratory (CZE)^34,35^, a geological setting characterized by alternating limestone and mudstone strata, resulting in varying hydrological connectivity within the carbonate-rock aquifers. Two footslope wells, located in thinner and less connected fractured mudstone-dominated aquifers, are more hydrologically isolated and contain old carbon sources (*e.g.*, > 4,500 years), while other wells in limestone-dominated aquifers having wider fractures exhibit greater hydrological connectivity due to minor karstification and higher permeabilities^5,36,37^. Previous studies have indicated the importance of connectivity through seepage transport of soil microorganisms into groundwater^15,16,17,18^. Consequently, the Hainich CZE well network enables comparisons of microbiome variability and stability across aquifers of different levels of hydrological connectivity.

Capitalizing on a decade-long groundwater survey comprising 815 bacterial 16S rRNA gene-sequence datasets and accompanying hydrochemical analyses, 226 DOM analyses, and 387 metabolomic analyses combined with 174 seepage microbiome sequencing datasets, this study provides the most comprehensive assessment of groundwater microbiome variability and stability, and their potential drivers to date. To better assess the responses of precious groundwater resources to disturbances caused by increasing hydroclimatic extremes, this investigation addresses four key unknowns: (i) the extent to which microbiomes of hydrologically connected vs. more isolated groundwater exhibit similar temporal dynamics and stability, (ii) the means by which assembly mechanisms explain microbiome variability, (iii) the extent to which seepage-associated microorganisms contribute to microbiome variability, and (iv) the means by which alterations in groundwater metabolome/DOM explain microbiome variability. By integrating microbial, hydrochemical, and metabolomic datasets, this study reveals critical insights into the responses of groundwater ecosystems to environmental disturbances and hydroclimatic extremes. These findings have broad implications for predicting groundwater resilience and informing sustainable management strategies under future climate scenarios.

## Results

### Hydrological seasonality and long-term variability of groundwater microbiomes

Our 10-year time series revealed distinct and persistent regular patterns of variation with alternating greater and lesser similarity over a 12-month period. While these variations are on a 12-month cycle, we use the descriptive term ‘sinusoidal’ because groundwater systems can integrate a number of potentially seasonally varying signals and lag times. These sinusoidal patterns in the similarity of shallow groundwater (< 100 m depth) microbial communities over time across all wells, despite differences in groundwater hydrochemical parameters and microbiome composition among wells (Fig. 1, Supplementary Fig. 1a, b). These patterns demonstrated consistent periodicity, with distinct peaks in similarities approximately every 12 months and minima roughly 6 months after each peak. This behavior was not confined to a single bacterial phylum, as approximately half of the bacterial phyla detected displayed sinusoidal patterns (Supplementary Tab. 1). Similar sinusoidal patterns were observed for hydrochemical properties of the groundwater (Supplementary Fig. 1c).

**Fig. 1.**
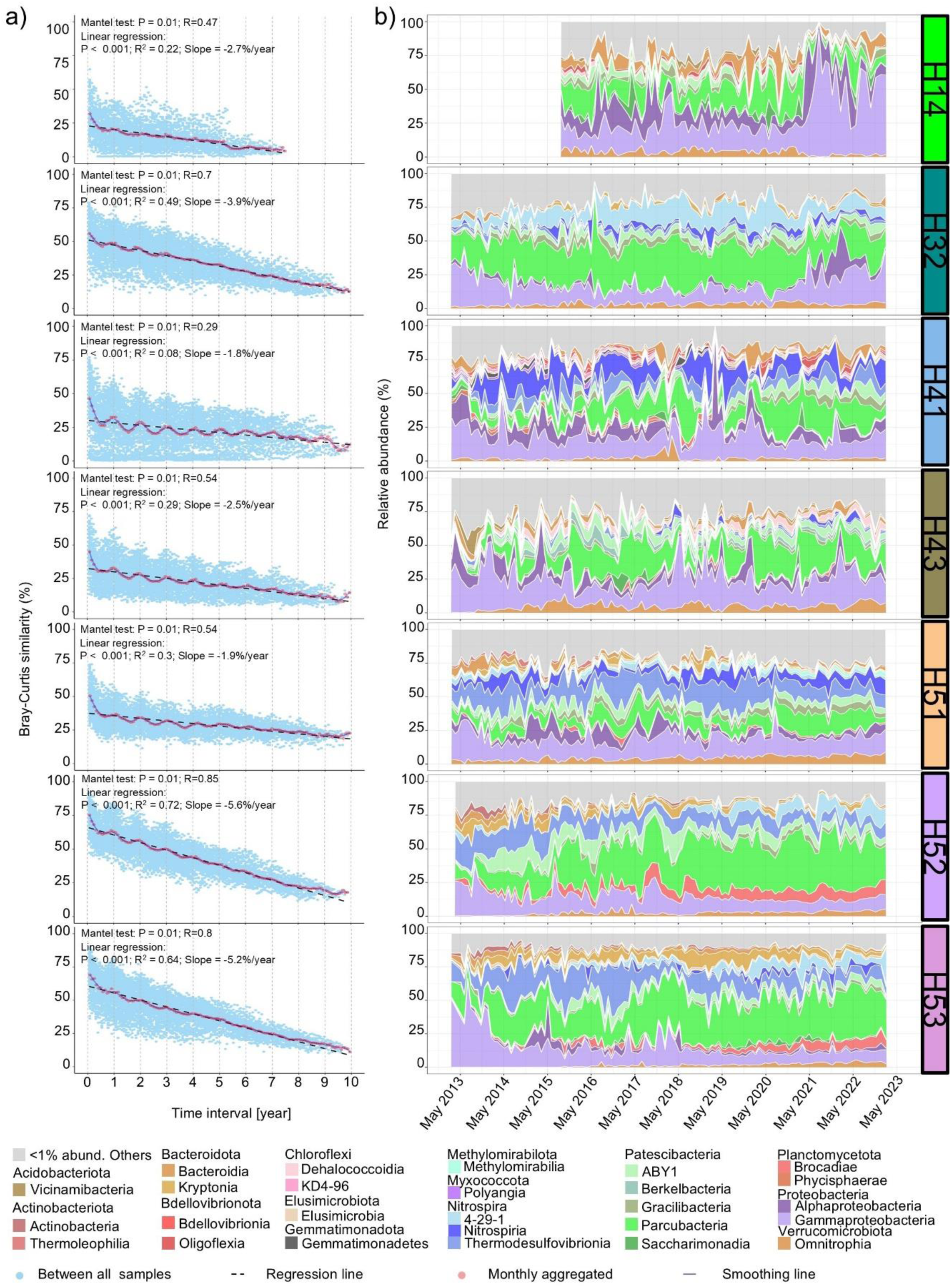
Sinusoidal and long-term variability in groundwater microbiomes. **a)** Temporal patterns in groundwater microbial community similarity: blue points represent the raw Bray-Curtis similarity between all samples against their sampling time intervals while red points represent the monthly average Bray-Curtis similarity between all samples. Regression lines and smoothing lines are applied to better envision temporal patterns. The turnover rate (%/year) is calculated as the slope of the regression lines. b) Temporal patterns in groundwater microbial community diversity at different wells: colors represent bacterial classes with relative abundances greater than 1%. The first six years of microbiome data at wells of H41, H43, and H52 were published in Yan *et al*^13^.

Sinusoidal amplitudes of microbiome community similarities varied among wells. Distance-based redundancy analysis (dbRDA) indicated that hydrological seasons (*i.e.*, early summer, later summer, early winter, and later winter) accounted for one to six percent of the microbiome variation (P < 0.05; Supplementary Fig. 2), except in well H53. In contrast, the phylogenetic structure of groundwater microbiomes exhibited weaker or no sinusoidal patterns (Supplementary Fig. 3). Only one to three percent of the variation in phylogenetic structure at four wells (H32, H41, H43, and H52) was influenced by hydrological seasons (dbRDA, P < 0.05). These findings suggest that various responses to changing hydrological seasons elicited by phylogenetically closely related microorganisms may offset seasonal effects.

Beyond sinusoidal patterns, all groundwater microbiomes showed long-term variability, as indicated by Mantel tests (P < 0.05; Fig. 1a). Microbial community similarity declined over time across all wells, with temporal turnover rates (*i.e.,* the rate of community similarity change) ranging from 1.8% to 5.6% per year (Fig. 1a). While the phylogenetic structures of the microbiome compositions also exhibited long-term variability (Mantel tests, P < 0.05), their temporal turnover rates were lower (0.8-2.6% per year, Supplementary Fig. 3). These results suggest that roughly half of the turnover in groundwater microbiomes may be directed towards phylogenetically closely related microorganisms.

### Contrasting temporal patterns of groundwater microbiomes

Groundwater microbiomes exhibited contrasting short-term variations, with mean Bray-Curtis distances between samples at one-month intervals ranging from 68% at well H14 to 25% at well H52, reflecting the strength of community composition fluctuations over time (Fig. 1b and 2a). In contrast, groundwater hydrochemical parameters, indicators of environmental changes, showed more consistent short-term variations, with mean Euclidean distances ranging from 14% to 26% (Supplementary Fig. 4). This disparity in short-term variations between hydrochemical parameters and microbiome compositions suggests varying levels of microbiome resistance to environmental changes, with greater short-term microbiome variation corresponding to lower resistance.

**Fig. 2.**
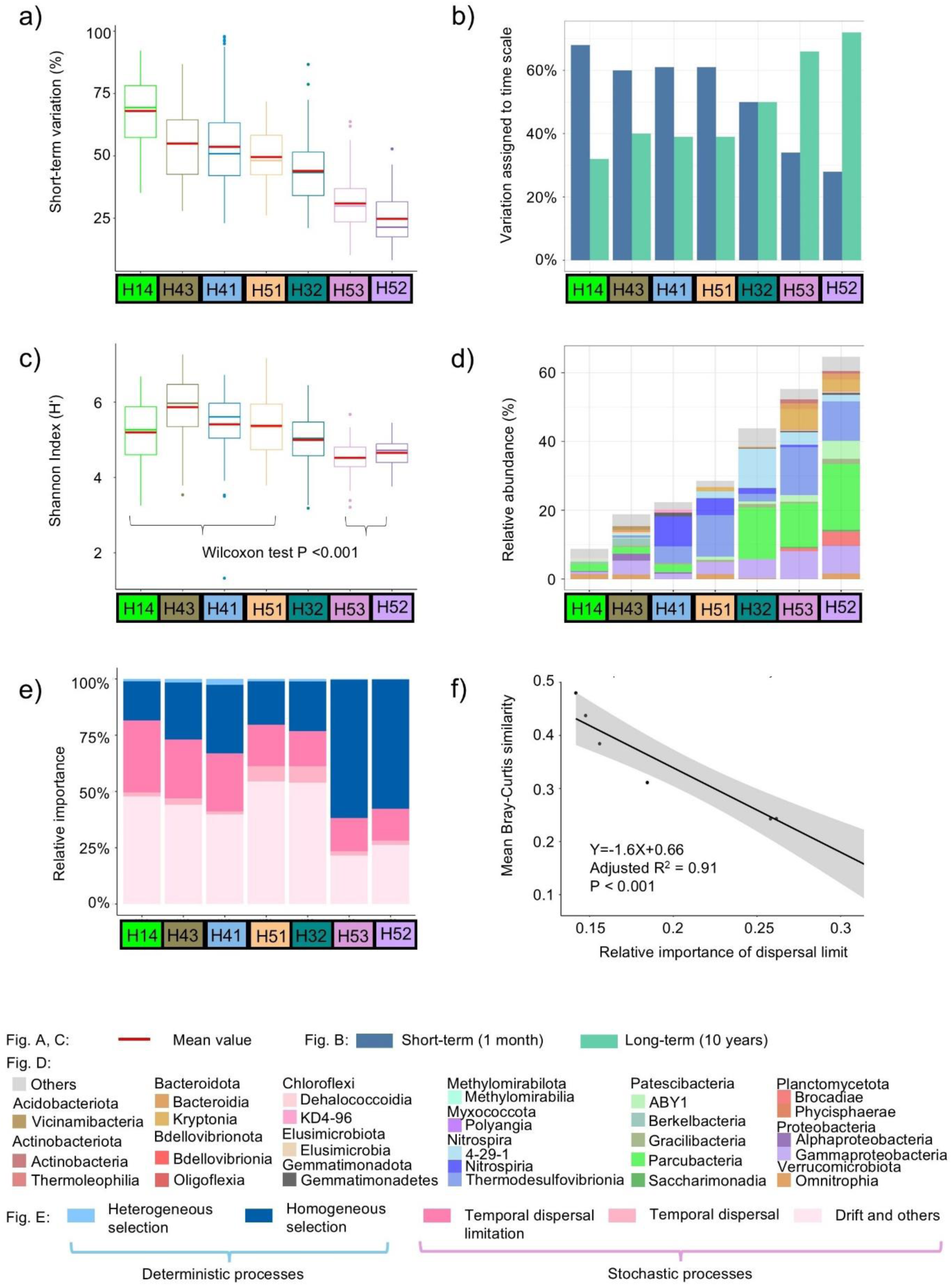
Distinct temporal patterns of groundwater microbiomes. **a)** Short-term variations in groundwater microbiome diversity based on Bray-Curtis distances between samples collected one month apart (short red horizontal lines denote corresponding mean values). b) Variation in groundwater microbiomes assigned to short-term (1 month in blue bar) and long-term (10 years in green bar) time scales. c) Shannon indices of groundwater microbiomes (short red horizontal lines denote corresponding mean values). d) Fraction of core microorganisms (ASVs present ≥ 80% of collected well samples; colors represent different bacterial classes). e) Relative importance of groundwater microbiome assembly processes: green hues represent deterministic processes (heterogeneous selection and homogeneous selection), while red hues represent stochastic processes (dispersal limitation, horizontal dispersal, and drift and others). Since these analyses were conducted on a single site over time, the dispersal here was related to time rather than space. f) Significant linear model between the relative abundance of (temporal) dispersal limitation and temporal stability of groundwater microbiome diversity (using mean Bray-Curtis similarity).

Pronounced short-term (1 month) variability was observed in microbiomes at wells in hydrologically connected aquifers (H14, H43, H41, and H51), while pronounced long-term (10 years) variability was detected in wells representing more hydrologically isolated groundwater (H52 and H53; Fig. 2b). Microbiomes with pronounced short-term variability yielded greater mean Shannon index values (5.2-5.9) and smaller fractions of core microorganisms (9-29%), compared to those with pronounced long-term variability, which exhibited lower Shannon indices (4.5-4.7) and larger core bacterial fractions (55-65%; Fig. 2c, d). Microbiomes at well H32 exhibited equally elevated short-term and long-term variability, with intermediate characteristics between hydrologically connected and isolated groundwaters, which may reflect fluctuating hydrological connectivity at this site (Fig. 2).

Microbiomes in hydrologically connected groundwater exhibited lower resistance but higher resilience, as indicated by their lower temporal turnover rates (1.8-3.9% year^-1^) compared to hydrologically isolated groundwater (5.2-5.6% year^-1^; Fig. 1a). Here, resilience refers to the ability of groundwater microbiomes to maintain community composition over time despite ongoing environmental change (as suggested by temporal variations in groundwater hydrochemical parameters across all wells; supplementary Fig. 1c). A low resilience is indicated by a high microbial community turnover rate. Temporal stability, measured by mean pairwise Bray-Curtis similarities, was significantly lower in microbiomes of more hydrologically connected groundwater (16-31%) than their hydrologically isolated counterparts (44-48%; Fig. 3a).

**Fig. 3.**
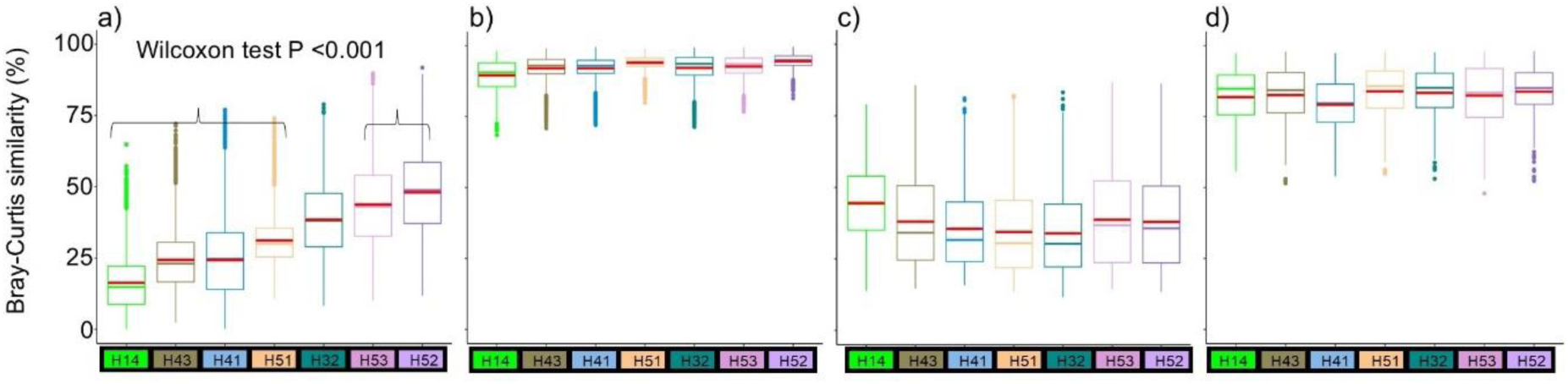
Temporal stability of a) groundwater microbiome compositions, b) metabolic potential of groundwater microbiomes, c) metabolome compositions, and d) bulk DOM compositions across wells per Bray-Curtis similarity. The metabolic potential of groundwater microbiomes in figure c) was inferred from 16S rRNA gene-sequences. Red horizontal lines denote mean values.

### Impact of seepage-associated microorganisms on groundwater microbiome variations

To appraise the extent of variability in groundwater microbiomes driven by soil-borne microorganisms transported with the seepage into the groundwater, we compared 16S rRNA gene datasets^16^ from soil seepage (23-60 cm depth) in local recharge areas with those from groundwater microbiomes. Seepage microbial diversity, dominated by Gammaproteobacteria (32%), Alphaproteobacteria (24%), and Parcubacteria (10%), differed significantly from that of groundwaters (Permutation tests, P =0.001; Supplementary Fig. 5).

SourceTracker analyses revealed that hydrologically connected groundwater harbored significantly higher fractions of seepage-associated microorganisms (2.4-8.9% of the total community) than did hydrologically isolated groundwaters (0.2-1.7%; Wilcoxon tests, P < 0.001; Fig. 4b). Despite these variations across wells, however, seepage-associated microorganisms consistently accounted for a mere four percent of groundwater microbiome variation (dbRDA; supplementary Fig. 2). These results suggest that low survival rates of seepage-associated microorganisms limit their contribution in shaping groundwater microbiome composition.

**Fig. 4.**
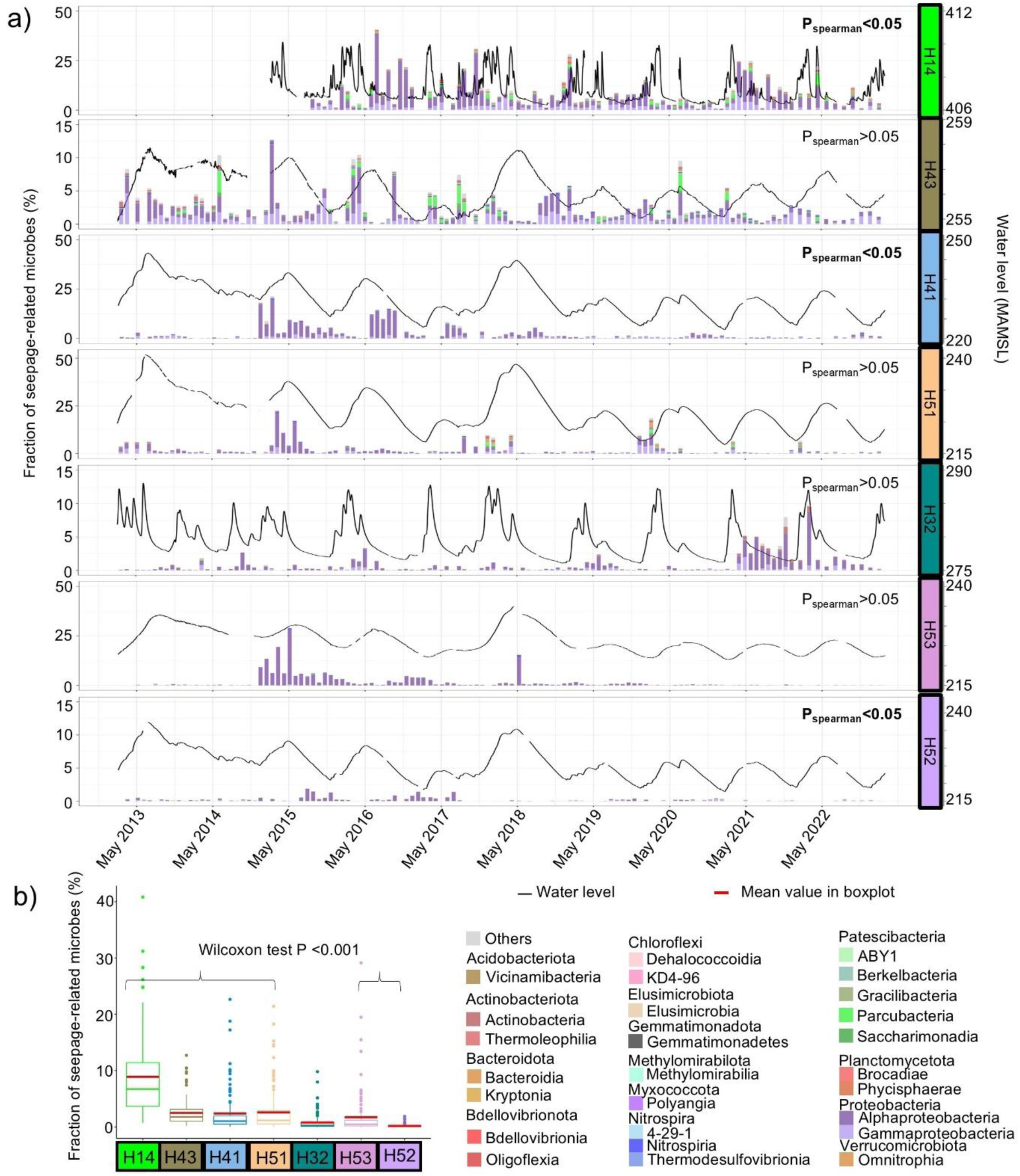
Fraction of seepage-associated microorganisms in groundwater as assessed by SourceTracker. **a)** Temporal patterns of seepage-associated microbial diversity: colors indicate microbial classes; black lines delineate groundwater level fluctuations; Spearman correlation assessed the relationships between seepage-associated microbe fractions and groundwater level fluctuations. b) Fraction of seepage-associated microorganisms across wells: red horizontal lines denote mean values.

A significant correlation was observed between the proportion of seepage-associated microorganisms and groundwater level fluctuations at wells H14 and H41 (Fig. 4), while a significant correlation between precipitation and groundwater levels was observed only at the shallowest well H14 (P_Pearson_ = 0.03; r_Pearson_ = 0.04; Supplementary Fig. 6). The fractions of seepage-associated microorganisms at wells H41 and H43 were significantly greater during groundwater recharge (rising water levels; 3.9% and 3.1%, respectively) than groundwater recession (falling water levels; 1.4% and 2%, respectively; Wilcoxon test, P < 0.01). Seepage-associated microbial diversity in groundwater was dominated by Alphaproteobacteria (mainly *Caulobacter*, *Sphingobium*), Gammaproteobacteria (mainly *Polaromonas*, *Rhodoferax*), and Parcubacteria (mainly *Candidatus* Adlerbacteria, *Candidatus* Nomurabacteria).

### Contrasting assembly processes in groundwater microbiomes

Microbiome composition was driven primarily by stochastic processes (stochasticity: 67-82%) in hydrologically connected groundwaters and deterministic processes in more hydrologically isolated groundwaters (stochasticity: 38-42%; Fig. 2e). Temporal dispersal limitation (dispersal limitation over time at one site) ranged from 18% to 32% in hydrologically connected groundwaters and 14% to 15% in more hydrologically isolated groundwaters (Fig. 2e), indicative of more dynamic population structures in the former. This was supported by higher Shannon indices, smaller fractions of core community members, and increased seepage-associated microbial inputs in hydrologically connected groundwaters (Figs. 2c, d; 4b). Correlations between elevated temporal dispersal limitations and lower temporal stabilities in groundwater microbiomes were confirmed by a linear regression model (Fig. 2f).

Results of dbRDA analyses highlighted the importance of environmental selection, particularly homogeneous selection, in driving microbiome assembly in hydrologically isolated groundwaters (Supplementary Tab. 2). Hydrochemical parameters accounted for 45-46% and 12-30% of microbiome variation in hydrologically isolated groundwaters and connected groundwaters, respectively. Groundwater level fluctuations yielded the greatest impacts, accounting for 14-21% and 25% of microbiome variation in hydrologically isolated and connected groundwaters, respectively (dbRDA, Supplementary Fig. 2). Regression analyses yielded significant long-term groundwater table declines across all wells, despite periodic recharge, with rates ranging from 7 cm year^-1^ at well H14 to 126 cm year^-1^ at well H41 (Supplementary Fig. 6). Other hydrochemical parameters (*e.g.,* temperature) also varied significantly with groundwater levels, leading to significant changes in groundwater environments over time (Mantel test, P < 0.05; Supplementary Fig. 1, 7).

Collectively, hydrochemical parameters, hydrological seasons and seepage-associated microbial input accounted for 15-33% and 47-50% of microbiome variation in hydrologically connected and isolated groundwaters, respectively (Supplementary Tab. 2).

### Functional potentials of groundwater microbiomes

PICRUSt2 identified 390 distinct MetaCyc metabolic pathways from 16S rRNA gene-sequence datasets. Much like the case for groundwater microbiome diversity, the potential metabolic pathway compositions exhibited both sinusoidal patterns and long-term variability (except well H41 which exhibited only sinusoidal patterns; Fig. 5a). Compared to microbiome diversity metrics, the potential metabolic pathway compositions exhibited higher temporal stability (mean Bray-Curtis similarity = 90-95%) and resilience (temporal turnover rates = 0.08-0.9% year^-1^; Fig. 2b and 5a).

**Fig. 5.**
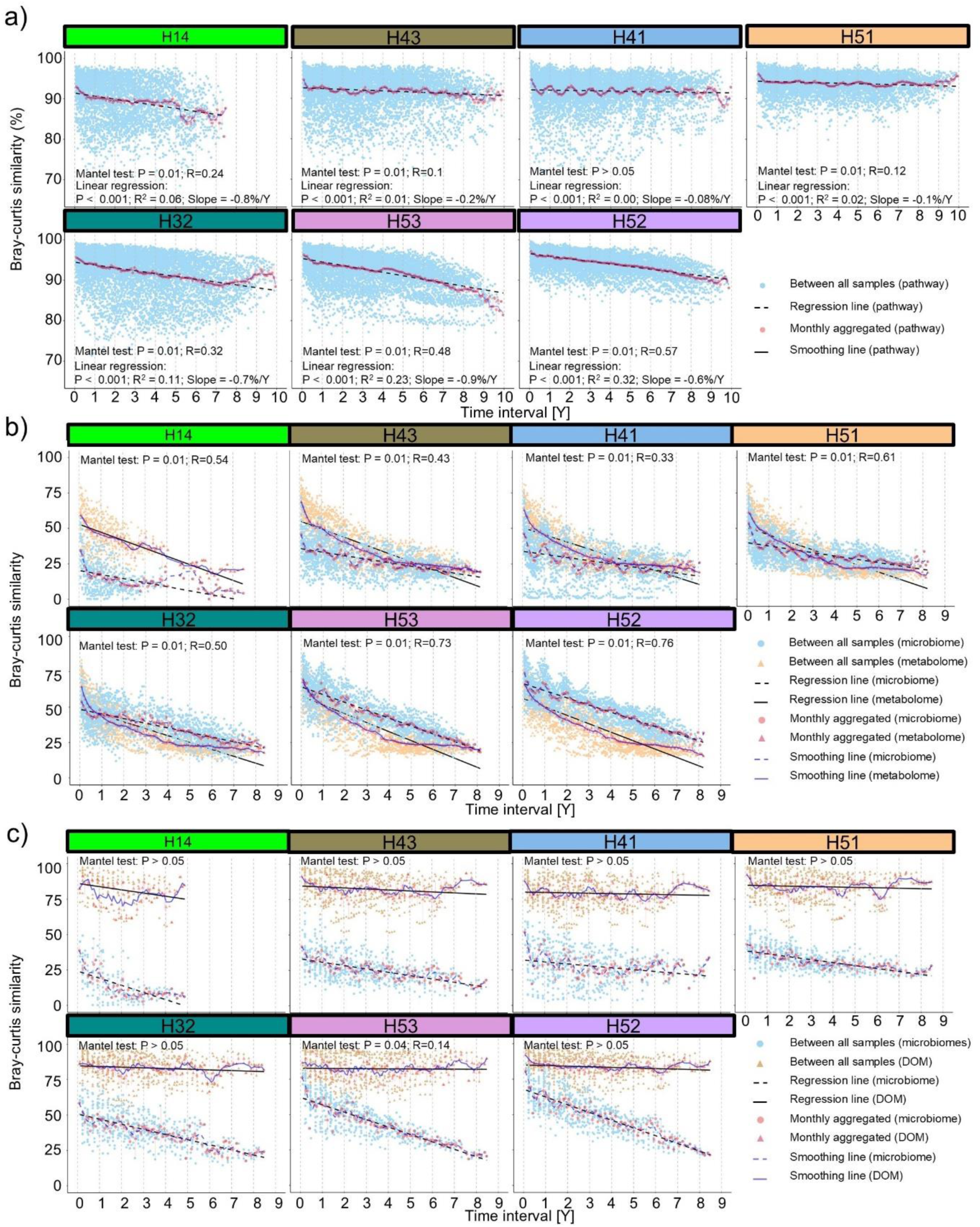
Temporal patterns of metabolic potential of microbiomes, metabolomes, and DOM compositions. **a)** Temporal patterns of metabolic pathways predicted by PICRUSt2 using 16S rRNA gene-sequences; b) Temporal patterns of groundwater metabolomes alongside a subset of concurrently sampled groundwater microbiomes; c) Temporal patterns of groundwater DOM compositions alongside a subset of concurrently sampled groundwater microbiomes. Mantel tests were conducted to assess correlations a) between changes in predicted pathway and time; b) between changes in microbiomes and metabolomes; c) between changes in microbiomes and DOM.

### Temporal patterns of groundwater metabolome compositions

LC-HRMS-derived metabolomics revealed both long-term variability and, albeit weak, sinusoidal patterns in the organic landscape (Fig. 5b). Significant correlations were observed between variations in microbiome and metabolic compositions across all wells (Mantel test; Fig. 5b). Variations in metabolomes, indicated by Shannon indices and primary axes of Principal Coordinates Analyses (PCoA), accounted for 11-16% and 39% of microbiome variation in hydrologically connected and isolated groundwaters, respectively (dbRDA; Supplementary Fig. 2).

Metabolome variation, together with hydrochemical parameters, hydrological seasons, and seepage-associated microbial input accounted for 31-59% of microbiome variation, roughly 10% more than when metabolome variation was excluded (dbRDA; Supplementary Tab. 2). In addition, the temporal stability of metabolomes (34-45%) was more consistent across wells than that of groundwater microbiomes (per mean Bray-Curtis similarity; Fig. 3).

### Temporal patterns of groundwater DOM concentration and compositions

Dissolved organic carbon (DOC) concentrations ranged from below detection limit (< 0.5 mg L^-1^) to 2.8 mg L^-1^, with higher concentrations detected in shallow wells H14, H43, and H32 (Supplementary Fig. 8). Variations in DOC accounted for one to six percent of the microbiome variation in wells H41 and H43, which were most affected by hydrological seasons (dbRDA; Supplementary Fig. 2). While bulk DOM compositions elucidated via DI-HRMS lacked significant correlation with microbiome variations (Mantel test; Fig. 5c), they exhibited greater resilience (lower turnover rates) and temporal stability (greater mean Bray-Curtis similarity: 79-84%; Fig. 3, 5c) than microbiomes.

To link DOM composition to microbial functions, we compared the compound classes inferred from DI-HRMS data to the potential degradation pathways inferred from 16S rRNA gene-sequences (Supplementary Tab. 3). The relative abundances of most degradation pathways were significantly greater in microbiomes in hydrologically connected than their hydrologically isolated counterparts (Wilcoxon test, P < 0.001), except for C1 compound utilization and polymeric compound degradation.

The relative abundances of two of the six DOM compound classes identified differed significantly between hydrologically connected and isolated groundwaters (Fig. 6; Supplementary Fig. 9). Condensed aromatic structures, likely derived from Muschelkalk sediments and underlying Permian/Jurassic formations, terrestrial plant material, or microbial necromass^38,39,40^, were more abundant in hydrologically connected groundwaters (0.15-0.4% vs. 0.04-0.08%; Wilcoxon test, P < 0.01; Fig. 5). This suggests that higher levels of condensed aromatics corresponded to an increased degradation potential for these compounds (Fig. 5). Conversely, relative abundances of peptide-like DOM, potentially sourced from extracellular enzymes and microbial necromass^39^, were significantly lower in hydrologically connected groundwaters (0.12-0.2% vs. 0.22-0.36%; Fig. 6). Yet, microbiomes in hydrologically connected groundwaters exhibited greater degradation potential for amino acids, even though their potential ability to synthesize them was not consistently lower (Fig. 6; Supplementary Fig. 10). This suggests that these communities recycle environmental proteins to a greater extent than their counterparts in more hydrologically isolated groundwater.

**Fig. 6.**
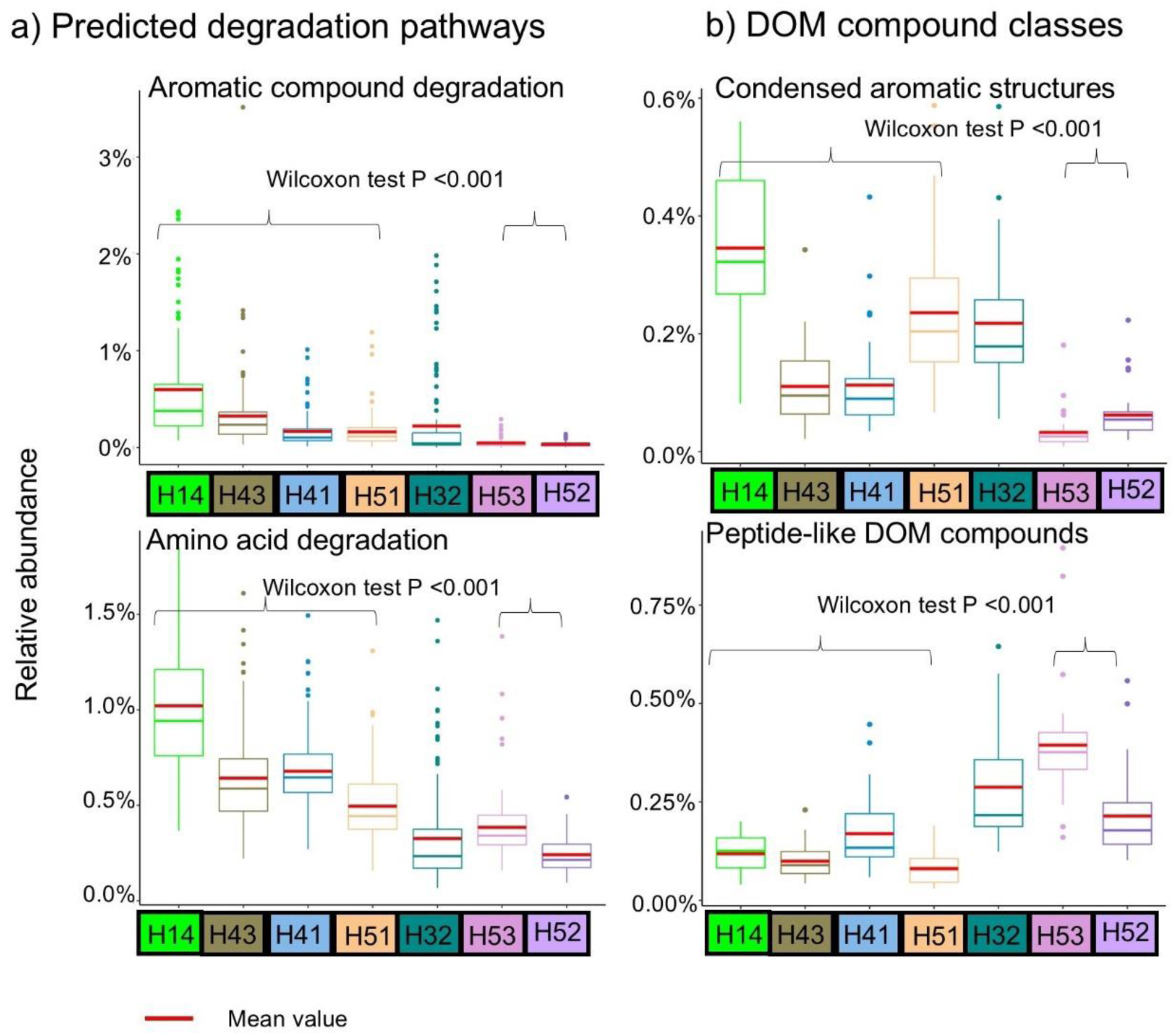
Boxplot of selected a) predicted degradation pathways and b) DOM compound classes. The predicted degradation pathways were inferred from 16S rRNA gene-sequences, while DOM compound classes were inferred from DI-HRMS derived bulk DOM. Red horizontal lines denote mean values.

Between May 2020 and October 2021, condensed aromatic structures and unsaturated hydrocarbons significantly increased at shallow wells H14 and H32 (and others). Mean relative abundances of condensed aromatics rose from 0.27% to 0.44% and from 0.19% to 0.39% (Wilcoxon test, P <0.01), while unsaturated hydrocarbons increased from 2.4% to 3.7% and from 2% to 3.8% at wells H14 and H32, respectively (Wilcoxon test, P <0.01). Following these increases, the degradation potential for aromatics and carboxylic acids at H14 and H32 rose significantly, beginning in March and April 2021, respectively. Aromatic degradation potential increased from 0.4% to 1.4% and from 0.06% to 1% (Wilcoxon test, P <0.001; Supplementary Fig. 11), while carboxylic acid degradation potential increased from 0.3% to 0.6% and from 0.07% to 0.3% at wells H14 and H32, respectively (Wilcoxon test, P <0.001; Supplementary Fig. 11). The relative abundance of Gammaproteobacteria rose significantly at H14 (30.4% to 69.6%) and H32 (15.7% to 34.7%; Wilcoxon test, P <0.001; Fig. 1b). These results suggest delayed changes in microbial composition and function in response to changes in DOM in shallow groundwater wells.

## Discussion

Our decade-long study of groundwater microbiomes unveiled hydrological seasonality and long-term variability, challenging the traditional view of groundwater microbiomes as static enteties^7,36,41^. Trends of continuous change in groundwater microbiomes were previously reported by Yan et al.^13^ from the same study sites, but sinusoidal patterns emerged only with this 10-year analysis. These patterns follow local seasonal hydrological changes, with hydrochemical parameters exhibiting annual sinusoidal patterns corresponding to groundwater level variations driven by meteoric recharge^5^ (Supplementary Fig. 1, 6, 7). Similar periodic groundwater level fluctuations have been observed in other shallow aquifers (< 100 m depth), especially in karst systems^10,11,12^. Sinusoidal patterns in microbiomes were never reported, however, even though shifts resulting from environmental changes were reported^12,22,42^, perhaps owing to shorter study periods and/or lower observation frequencies. Annual sinusoidal patterns, driven by seasonal factors (*e.g*., day length, temperature, nutrient availability), are common in long-term studies of marine and lake environments, particularly marine surface layers^25,26,43,44^.

Sinusoidal patterns in subsurface microbiome diversity may arise from specific microorganisms being periodically favored by fluctuating environmental conditions, such as redox potential and/or groundwater levels^11,13^. Microorganisms benefitting from high groundwater levels include those remobilized from rock surfaces in the vadose zone, as local carbonate rocks and planktonic groundwater communities share up to 40% species diversity^45^. This fraction might be even higher in porous aquifers^46^, rendering them important seeding banks for groundwater microbiomes^21^. In addition to enhancing microbiome stability by increasing homogeneous selection (*e.g*., well H41; Fig. 2e), the sinusoidal patterns characterized in this study drastically improve predictions of microbiome change^47,48^.

The Hainich CZE provides an ideal setting to study microbiome stability in carbonate-rock aquifers of varying hydrological connectivity. Our extended in-depth characterization of seven selected wells considered several distinct microbiomes along the groundwater monitoring transect^49^, revealing contrasting temporal patterns and stability in groundwater microbiomes over a mere 6 km. The microbiomes of more isolated groundwaters with high temporal stability (*e.g*., wells H52 and H53) are influenced largely by deterministic processes, while those in hydrologically connected groundwaters are shaped by stochastic processes (and as such exhibit lower temporal stability). In well H32, an intermediate aquifer system, groundwater microbiomes exhibited both elevated short- and long-term variability, reflecting sporadic hydrological connectivity likely resulting from complex flows in fractured sedimentary bedrock aquifers^5^.

Our results corroborate the findings of recent groundwater studies in other geological settings^11,50,51^, suggesting that hydrological connectivity reduces the temporal stability of groundwater microbiomes by increasing the role of temporal dispersal limitation via microbial immigration. Thus, hydrological connectivity lowers the resistance of the groundwater microbiome while increasing its resilience. Furthermore, microbial immigration during recurrent groundwater recharge events represents intermittent disturbances that promote resilience and create opportunities for coexistence by preventing competitive exclusion, thereby promoting diversity^52^.

In near-surface aquifers, seepage-associated microbial input is a key contributor of microbial immigration, increasing the importance of temporal dispersal limitation in microbiome assembly^20,22^. Our findings suggest that 0.2-8.9% of the total groundwater microbiome is seepage-borne, likely derived during periodic groundwater recharge or single hydrological extreme events (particularly relevant in karst systems). Recurrent immigration events over thousands of years may have enhanced the persistence of these invaders in the groundwater microbiome, as up to 12.5% of groundwater core ASVs have been identified as being seepage-associated. As these invasive taxa survive and partake in community coalescence, they compete with resident taxa for resources, and closely related species compete for similar resources^53^. Such competition between species oftentimes results in divergence in niches, including increased host specificity of episymbiotic microorgansisms (*e.g*., Patescibacteria), to reduce competition costs^54^.

The effects of hydrological connectivity on microbiome stability extends beyond groundwater, with similar patterns reported in oceans and other aquatic ecosystems^55,56,57,58,59,60^. Deep ocean microbiomes (890 m depth) are more temporally stable, with mean Bray-Curtis similarities around 64%, compared to 40% in the more hydrologically connected surface layers (5-20 m depth) where currents and nutrient mixing enhance microbial exchange^55,60^. Somewhat surprisingly, surface/shallow ocean microbiomes are more stable than those of hydrologically connected groundwaters, and show reduced long-term community turnover^43,55,60^. Compared to groundwaters, marine microbiomes exhibit higher resilience (*i.e.*, lower turnover rates) throughout all depths/layers^43,55,60^. This suggests that groundwater systems are more vulnerable and less resilient than marine systems^61^.

The functional potentials of all seven groundwater microbiomes exhibited remarkable temporal stability and resilience despite notable variations in community composition. This aligns with previous groundwater studies that report high functional redundancy in these ecosystems^11^. Functional redundancy, *i.e.*, multiple microbial taxa performing overlapping ecological roles, ensures ecosystem stability amid species turnover^62^. The observed resiliencies in functional potential suggests that, despite environmental fluctuations, these microbial communities retain the capacity to adapt while maintaining their functional integrity^8,21^. Significant correlations were observed between microbiome and metabolome changes, pointing to a form of functional redundancy achieved through metabolic diversity. Such a mechanism enables different microbial taxa to produce varied metabolites that fulfill similar ecological functions^63^. Metabolome compositions exhibited greater temporal consistency across wells than microbiomes. This stands to reason as not all detected metabolites are microorganism-related, which in and of itself highlights the influence of additional controlling factors^28,30^. The lower temporal stability of groundwater metabolomes compared to corresponding microbiomes’ elevated functional potentials corroborates earlier findings which reported substantial metabolome variability at the same site^29^.

Distinct temporal patterns were observed in groundwater metabolome and DOM compositions, largely attributable to differences in analytical methods that preferentially target different compound classes^14,29^,. The high temporal stability of DOM across wells may result from its composition, which consists of lignin-degradation products and polyphenolic leachates from plant matter in topsoil^36,64,65^ (Supplementary Fig. 9). These compounds resist microbial degradation, contributing to their persistence in the environment^36,64,65^. Despite comprising less than 5% of total DOM, the microorganism-associated DOM fraction likely plays a significant role in subsurface organic matter cycling. Its influence on spatial and temporal variation in microbiome composition and function^36,66,67^ was particularly evident in the shallowest well (*e.g.*, H14, H32). The contrasting dynamics observed between DOM and metabolomes emphasize the value of integrating diverse analytical approaches^28,32^.

This long-tern study shows that contrasting hydrogeological conditions render the Hainich CZE an ideal setting to elucidate the various mechanisms driving groundwater microbiome stability and vulnerability in carbonate-rock aquifers. Highly karstified regions, rich with extensive fractures and conduits, are globally crucial for drinking water as they facilitate rapid groundwater flow and abundant water storage (in contrast to less permeable, dense carbonate rocks)^4^. Karst aquifer microbiomes are typically influenced most largely by stochastic processes and are more susceptible to disturbances from surface-derived inputs (*e.g*., organic contaminants, pathogens)^22,67,68,69^. As such, improper surface management can rapidly increase their vulnerability and threaten groundwater health. Extensive source control to limit surface contaminant ingress to aquifers is crucial for maintaining the health of these ecosystems^23^. Moreover, hydroclimatic extremes, such as heavy precipitation and drought, evidently exacerbate groundwater vulnerability. These extremes facilitate the high ingression of surface-derived organic molecules (*e.g.*, xenobiotic substances) into groundwater by evading microbial processing^14^. The high ingression of surface-associated bacteria during recharge and heavy precipitation events raises concerns about pathogen contamination, threatening groundwater quality. These findings underscore the vulnerability of highly hydrologically connected aquifers, such as karst aquifers, under hydroclimatic extremes.

In contrast, microbiomes in more hydrologically isolated groundwaters (*e.g.*, dense limestone formations) tend to be governed by deterministic processes and often resist disturbances^70,71^. However, this study demonstrates that continuous environmental changes result in decreasing temporal stability in microbiomes in hydrologically isolated groundwater over time, even in the absence of contamination. Such changes may be driven by prolonged droughts affecting groundwater recharge. While these systems may exhibit low short-term variability and greater resistance to contamination, their recovery following severe contamination is likely prolonged, as a new ecological balance emerges.

## Conclusion

Overall, the results of this study show that both hydrologically connected and isolated groundwater ecosystems in carbonate aquifers are increasingly at risk under scenarios with more frequent hydrological extremes that alter the connections between surface water and groundwater. As hydroclimatic extremes intensify, the persistence of core microbial functions becomes increasingly important to maintaining ecosystem stability. The findings discussed here underscore the importance of innovative integrated management strategies (*e.g*., robust surface control measures, long-term monitoring of aquifer health) to safeguard Europe’s groundwater resources from the stresses of contamination and climate change.

## Methods

### Study sites and sampling

The study site is located in the Hainich CZE in central Germany. Detailed site information and sampling procedures can be found in Kohlhepp *et al*.^34^, Küsel *et al*.^35^, and Lehmann and Totsche^5^. Briefly, the CZE features alternating thin–bedded limestone–mudstone in hillslope terrains, which is a common and widely distributed geological setting. The bedrock of the low–mountain hillslope in the eastern Hainich CZE consists primarily of Upper Muschelkalk of marine origin (Germanic Triassic), parts of which harbor abundant groundwater resources. Along this hillslope, our seven monitoring wells are embedded in a transect spanning 5.4 km (Supplementary Fig. 1A), encompassing various relief positions, aquifers, and depths^35^. Aquifer compartments at Hainich CZE differ significantly in oxygen availability. Wells H14, H32, H41, and H51 are oxic, H43 is suboxic (< 1 mg/L dissolved oxygen), and H52 and H53 are anoxic (< 0.1 mg/L dissolved oxygen). Redox potentials range around 400 mV for wells H14, H32, H41, and H51, and around 200 mV for wells H43, H52, and H53. The wells are situated in areas of extensive land management, including forest, pasture, and cropland. The shallowest well (H14), located on the upper hillslope, is the only well located in the preferential groundwater recharge area.

Groundwater samples were collected monthly to monitor standard hydrochemical parameters, microbial communities, and untargeted metabolomics. DOM was measured every three months. From February 2013 to February 2023, we collected 815 groundwater samples from seven groundwater wells for microbial analysis and hydrochemical parameters, 226 groundwater samples for DOM, and 387 samples for metabolomic analyses (Supplementary Fig. 12). Once the physico–chemical parameters of pumped groundwater stabilized, groundwater was collected from each well using a submersible pump (MPI, Grundfos) and placed into sterilized bottles. These samples included 5–10 L for microbiological analysis, duplicate 10 L samples for DOM analysis, triplicate 5 L samples for metabolomics, and 100 mL for DOC concentration measurements. Groundwaters from which to isolate genomic DNA was processed through 0.2 µm filters (PES or polycarbonate from Supor, Pall Corporation and Merck– Millipore, respectively) via vacuum pumping, and filters were stored at –80°C prior to DNA extraction. Acidified filtered groundwaters (0.7 µm filters, pH = 2 with HCl) were stored at 4°C in the dark until further DOM processing, while raw groundwaters were stored in the dark and chilled until further metabolomics analyses. Other groundwaters were filtered through 0.7 µm filters and stored at 4°C in the dark until further DOC quantification.

Groundwater hydrochemical parameters including groundwater levels, temperature, pH, specific electrical conductivity (EC_25_; reference T: 25°C), dissolved oxygen content, redox potential (ORP), acidity (neutralizing capacity), alkalinity (neutralizing capacity), total inorganic carbon (TIC), and ion concentrations were measured as described by Lehmann and Totsche^5^ and Kohlhepp *et al.*^34^. Element concentration including Ca, K, Mg, Na, and S were measured with ICP-MS (inductively coupled plasma mass spectrometry; 8900 Triple Quadrupole ICP-MS, Agilent, Germany), while major anions Cl^-^ was measured by IC (ion chromatography; Dionex IC20, Thermo Fisher Scientific, USA). DOC concentration was quantified as non-purgeable organic carbon on a vario TOC cube (Elementar Analysensysteme, Germany) with a detection limit of 0.5 mg L^-1^.

The sampling details for the 174 seepage samples collected in this study can be found in Hermann *et al*.^16^ and Lehmann *et al*.^72^ (Supplementary Fig. 1; Supplementary Tab. 4). Seepage sites were sampled regularly (biweekly) and on an event-basis (weekly). Seepage volumes ranging from 100 to 500 mL were filtered through 0.2 µm filters (PES; Supor, Pall Corporation) using a vacuum pump, and filters were stored at –80°C prior to DNA extraction.

### DNA extraction and amplicon sequencing

Groundwater and seepage genomic DNA were extracted using the DNeasy PowerSoil Pro Kit (Qiagen, Hilden, Germany) per manufacturer’s instructions, and extractions were stored at –20°C prior to PCR amplification. PCR amplification of bacterial 16S rRNA gene (V3–V4 region) was performed on an Illumina Miseq platform using v3 chemistry with primers Bakt_0341F and Bakt_0785R^73^. Most samples (see list in Supplementary Tab. 5) were sequenced in–house following the two–step PCR library preparation procedures described by Krüger *et al*.^17^. Amplicon libraries for some samples were generated using the NEBNext Ultra DNA Library Prep Kit for Illumina (New England Biolabs, MA), following methods detailed by Kumar *et al*.^74^ All remaining samples were processed at LGC Genomics (Berlin, Germany), as previously described^75^.

### Molecular composition of DOM

A detailed description of the methods used to elucidate the molecular compositions of DOM is given in Schroeter *et al.*^14^. Briefly, DOM was extracted from acidified filtered groundwater via solid phase extraction with PPL (styrene–divinylbenzene polymer) Bond Elut cartridges (Agilent Technologies) following the protocols of Dittmar *et al*.^76^. PPL extracts were kept at −80°C prior to DI-HRMS analyses. DI-HRMS analyses were conducted on an Orbitrap Elite mass spectrometer (Thermo Fisher Scientific, USA) with a mass resolution of 555,000 ± 9,000 at m/z = 251. The electrospray ionization (ESI) was run in negative mode with an ESI needle voltage of 2.65 kV. For each sample, 100 scans of m/z 100-1000 were acquired and averaged. Quality controls and molecular formula assignments were processed using DOMAssignR (https://github.com/simonschroeter/DOMAssignR). Spectra were further normalized to sum all peak intensities. We focused on DOM compound classes most likely to serve as substrates for microorganisms, including carbohydrates, condensed aromatic structures, lignins, lipids, peptide-like compounds, and unsaturated hydrocarbons. Identification of DOM clusters was based on their elemental N content and hydrogen/carbon and/or oxygen/carbon elemental ratios (Supplementary Tab. 6)^77^.

### Untargeted metabolomics

Sample extraction and analyses are described in detail in Zerfaß *et al*.^29^. Briefly, 5 L of filtered (GF/C, 1.5 µm, VWR) groundwater was subjected to solid phase extraction (SPE) in Strata–X 33 µm polymeric reversed phase cartridges (Phenomenex). Eluates (1:1 methanol:acetonitril) were dried (vacuum, nitrogen stream), and the organic residue was re-dissolved in 100 µl of 1:1 THF:methanol. Extracts (1 µl) were then analyzed by LC–HRMS (liquid chromatography–high-resolution mass spectrometry) on a Dionex UltiMate 3000 chromatography system coupled to a Q–Exactive Plus orbitrap mass spectrometer (Thermo Fisher Scientific), m/z-range 100-1,500, alternating acquisition in positive and negative mode. For this study, only positive-mode data was extracted.

To assure system suitability in the long-term sampling experiment, the LC-HRMS system was maintained with a weekly MS source cleaning, mass calibration, and consistency-check by injection of a standard (containing Fluorophenylalanine, P-Fluorobenzoic acid, Decanoic acid D-19) for which retention times and peak apex intensities were recorded. All samples were taken in environmental replicates and injected in triplicate analytical replicates, and each set of analytical replicates was processed in randomized sequence. Data processing for peak picking and feature assignment was carried out in XCMS as described in the stated reference. For Bray-Curtis similarity tests (details in succeeding section), replicate means were calculated and peak areas were normalized by the sum of all feature peak areas.

### Bioinformatics and statistical analyses

Most bioinformatics and/or statistical analyses were conducted in R version 4.2.2^78^ at a significance level of α = 0.05. After correcting the orientation of mixed-orientation reads and removing primers with Cutadapt^79^ (v 4.1), R package “dada2”^80^ (v1.26) was employed for quality filtering, denoising, inferring amplicon sequence variants (ASVs), and removing chimeras. Reads were truncated to 265 bp (forward reads) or 235 bp (reverse reads), excluding those with more than two expected errors and those truncated when the quality score was equal to or less than two. ASVs were generated by applying the DADA2 core algorithm and combining forward and reverse reads, and those that could not be aligned to the SILVA reference database^81^ (v138.1) using Mothur (v1.46.1) were removed. After removing chimeric sequences, taxonomy was assigned to the remaining ASVs based on the SILVA taxonomy reference database v138.1. Further downstream sequence analysis was conducted using the R package “phyloseq”^82^ (v1.42), and a phylogenetic tree was constructed using FastTree^83^ (v2.1) after aligning genes with Muscle^84^ (v5) and trimming the alignment with trimAl^85^ (v1.4).

Recharge and recession phases (Supplementary Fig. 6 and Tab. 5) were defined by observed changes in groundwater levels. The recession phase was characterized by a sustained decline in groundwater levels for more than five consecutive days, ending in a minimum level or (for well H14) a significant rise to or above the maximum level for that phase. The recharge phase was defined in an opposite manner. We considered one hydrological year (between 1 May and 30 Apr) comprising four seasons: hydrological early summer (May to Jul), late summer (Aug to Oct), early winter (Nov to Jan), and late winter (Feb to Apr). To visualize the heterogeneity of groundwater environments, principal component analysis (PCA) was conducted with 15 hydrochemical parameters, including water level, water temperature, specific electrical conductivity, dissolved oxygen concentration, redox potential, acidity, alkalinity, total inorganic carbon, element and ion concentration (Cl^-^, S, Ca, K, Na, and Mg). Pearson and Spearman correlations from Hmisc (Rpackage v5.1-1) were used to determine significant correlations.

We used mean pairwise Bray-Curtis similarity to evaluate the temporal stability of groundwater microbiome composition and function, while short-term variabilities was assessed using Bray-Curtis distances for sample pairs from the same well (sampling intervals of 15 to 44 days). To explore temporal patterns, we first plotted raw pairwise Bray-Curtis similarity against sampling time intervals. To illustrate community similarity decay rates (resilience), we modeled the regression line of the raw pairwise Bray-Curtis similarity. To identify periodic patterns, we divided the pairwise Bray-Curtis similarity into monthly intervals (*e.g*., a time interval of 1 month: 15-44 days; a time interval of 2 months: 45-74 days), and applied a smoothing line based on a moving average filter. The same technique was applied to groundwater DOM and metabolomes using Bray-Curtis similarity, groundwater microbiome compositions at the phylogenetic level using UniFrac similarity, and environmental parameters using Euclidean similarity. The Bray-Curtis similarity (distances) and UniFrac similarity (distance) mentioned in this manuscript were based on relative abundance, while the Euclidean similarity (distance) was calculated after normalizing the aforementioned 15 hydrochemical parameters to exclude the effect(s) of absolute abundance (values) differences. Variation assigned to short-term or long-term variability is represented as the proportion of the respective variability in one-decade variability. We first evaluated the 10-year variability of microbiome compositions using the regression model of raw pairwise Bray-Curtis similarity, at which time long-term variability is then calculated by subtracting the aforementioned short-term variability from the 10-year variability.

To elucidate the impact of surface-subsurface connectivity in the temporal stability of groundwater microbial communities, we tracked changes in contribution of seepage-associated microorganisms to the groundwater microbiome via SourceTracker^86^ (Rpackage v1.0.1 under R version 4.0.0). During these analyses, we applied all seepage samples as the source and the groundwater samples as the sink. Analyses were performed on rarefied ASV (rarefaction depth of 1,000 reads) abundances using default settings (α = 0.001, β1 and β2: 0.01).

In this study, we define core species as bacterial ASVs present in > 80% of the groundwater samples collected in every well. Distance-based redundancy analysis (dbRDA) based on Bray-Curtis dissimilarity was carried out to assess and validate whether selected environmental parameters (*e.g*., hydrological seasons, incidence of seepage-associated microorganisms) significantly impact groundwater microbiome composition (Rpackage vegan; adjusted R^2^ was used). The significance of dbRDA tests was reported by permutation tests of “anova.cca”. The metabolic functions of groundwater microbial communities were predicted by PICRUSt2 software based on taxonomy annotations from 16S rRNA gene sequences^33^. We employed inferred community assembly mechanisms using a phylogenetic bin-based null model (iCAMP, Rpackage v1.6.5) to evaluate the contribution of ecological processes on groundwater microbiome assembly^87^. All analyses were performed using recommended default settings with 48 bins and confidence for null model significant tests.

## Data and code availability

Raw amplicon sequencing data reads for all studied samples have been deposited in the European Nucleotide Archive (details in the Supplementary Material). Raw DOM data from DI-HRMS were deposited under https://doi.org/10.17617/3.2TZM6C, while raw metabolome data from LC-HRMS were deposited via the Metabolights repository^88^ under MTBLS3450, MTBLS8433, and MTBLS11375. Groundwater hydrochemical parameters are provided as Supplementary Material. Raw amplicon sequencing data and codes will be released upon publication of the manuscript.

## Acknowledgements

This study is part of the Collaborative Research Centre AquaDiva of the Friedrich Schiller University Jena, funded by the Deutsche Forschungsgemeinschaft (DFG, German Research Foundation) – Project-ID 218627073 – SFB 1076. The authors would like to thank Heiko Minkmar, Falko Gutmann, René Maskos, and Stefan Riedel for groundwater sampling and on-site measurements/sample preparation, and Karel Castro Morales for helpful suggestions. Special thanks are extended to Anke Hädrich and Maria Fabisch for scientific coordination. KK, GP and KUT gratefully acknowledge support from the DFG under Germany’s Excellence Strategy – Project–ID 390713860 – EXC 2051.

## Author contributions

HW and KK conceptualized the manuscript. KK, MH, and HW provided microbiome data. SAS, AI, GG, and SET provided DOC and bulk DOM data. CZ and GP provided metabolomics data. RL, KL, and KUT managed field installations and provided hydrochemical parameter analyses. HW processed data, performed bioinformatic analyses and wrote the manuscript. All authors discussed the results and implications and commented on the manuscript at all stages.

## Competing interests

The authors declare no competing interests.

## Notes

### Competing Interest Statement

The authors have declared no competing interest.

